# Historical disturbances to rivers appear to constrain contemporary distribution of a river-dependent frog

**DOI:** 10.1101/2021.01.22.427759

**Authors:** Mark A. Linnell, Raymond J. Davis

## Abstract

Frogs dependent on lotic environments are sensitive to disturbances that alter the hydrology (e.g., water impoundments), substrate (e.g., debris torrents), and riparian vegetation (e.g., wildfires) of river ecosystems. Although rivers are often very dynamic, disturbances can push environmental baselines outside of narrowly defined ecological tolerances under which a species evolved. Short-lived lotic-dependent organisms, restricted to movements within the water or the riparian corridor, are at risk of local extirpations owing to such disturbances if they fragment and isolate affected populations from recolonizing source populations. In Oregon, USA, the foothill yellow-legged frog (*Rana boylii*) is at its northernmost range margin and has experienced an approximately 41% range contraction compared to their historical distribution. To inform conservation and management, we used species distribution models to identify environmentally suitable watersheds based on intrinsic baseline environmental variables, and then examined potential effects of human-caused alterations to rivers, including splash dams used to ferry timber downstream prior to 1957, large water impoundments, and adjacency to agricultural croplands. We used machine-learning in program Maxent and three different river layers that varied in extent and location of mapped rivers but contained distinct information to produce species distribution models which we then combined into a single ensemble model. Stream order, annual precipitation, and precipitation frequency were the highest ranked baseline environmental variables in most models. Watersheds with highly suitable baseline conditions in our ensemble model were negatively correlated with anthropogenic disturbances to rivers. Foothill yellow-legged frogs appeared to be sensitive to human-caused disturbances to rivers, perhaps indicative of their narrow ecological tolerance to in-river conditions. We do not anticipate variables in our model to change much through time. Rather, for conservation we identified potential legacy (spash dams) and ongoing human-caused disturbances that are more likely to change conditions for the species in the short- and long-term.

## Introduction

Lotic-dependent amphibians are frequently of conservation concern as they often have narrowly defined ecological tolerances and are sensitive to human-caused disruptions to hydrological processes, including alterations to substrate and flow, temperature, as well as disease, invasive species, and other factors (1,2). Identifying the range of conditions in which these amphibians can occur but also anthropogenic disturbances to rivers or competitive interactions that may have constrained their current distribution can inform species management and conservation, particularly at range margins (3). Species distribution models can be a key first step to identifying these factors.

Foothill yellow-legged frogs (*Rana boylii*) in western Oregon are at their northern range edge, have experienced an approximately 41% distributional contraction, and are currently being considered for protection under the United States endangered species act (4). Across their range, they are a species of conservation concern, and water development and diversions, including for agriculture, that cause inconsistent flow regimes, change average or variation in river temperature, water chemistry, and alter river-bed geomorphology, or provide conditions favorable to invasive species are thought to be the primary cause of observed declines (1,2,5,6). Foothill yellow-legged frogs inhabit and breed in slow-moving rivers that contain coarse substrates (e.g. gravel bars, larger cobbles, and bedrock >2 mm; (7)) with limited streamside vegetation in western Oregon and California (2). Breeding coincides narrowly with summer drought and warming river temperatures, and foothill yellow-legged frogs may be particularly sensitive to alterations to water temperature, and high flow volumes that wipe away egg-masses during breeding season (2,8–10).

River ecosystems in western Oregon have been altered by extensive human-caused disturbances in the 20^th^ and 21^st^ centuries, including extensive splash dams used to ferry timber downstream in mountainous areas, settlement and agriculture in valleys, and water impoundments (11,12). Splash dams, historically used to ferry timber downstream in western Oregon ((13); 1880 to 1956), and described as a ‘sustained pulse disturbance’, channelized and scoured river channels, and among other impacts, potentially reduced availability of substrates (e.g., coarse sand, pebbles, and cobbles) that foothill yellow-legged frog use for breeding and protection from predators. Impacts to the in-river environment from splash dams were thought to be broad and long-lasting (11,14,15). Settlement and agriculture, particularly in the Willamette Valley at the northernmost historical locations of frogs, precipitated widespread alterations to riverine and terrestrial environments that included a ~25% decline in length of the Willamette River (12). Cessation of annual burning by native peoples to clear vegetation increased recruitment of streamside vegetation, potentially shading rivers and reducing suitability for foothill yellow-legged frogs (12). Dams, although not as widespread in western Oregon as in other parts of their range, have been linked with direct negative impacts by altering river temperatures, flow volume, and timing, and indirect effects if those alterations facilitate invasion by non-native species (1,2). These disturbances are likely to negatively affect local, breeding populations of foothill yellow-legged frogs and potentially disrupt connectivity among populations via barriers and by increasing distances between populations (2).

Multimodel ensembles, by combining several models, can increase confidence in species distribution model predictions but also indicate uncertainty in prediction (16). River networks are frequently difficult to consistently map due to subtle differences in model algorithms, interpretation of topographic features and extent of network. Models that are unable to be reconciled *a priori* due to spatial scale, extent, or location inconsistencies of input spatial environmental predictor layers but that provide unique information may benefit from an ensemble model approach.

Our objectives were to build an ensemble of species distribution models for the foothill yellow-legged frog and to summarize human-caused disturbances to rivers, particularly in areas where the species historically occurred, based on suitable intrinsic environmental conditions, but now appears extirpated. We contrast our results with a previous model (4), summarize disturbances to rivers in western Oregon, and identify watersheds where models agree. Finally, we combined new species distribution models into a final ensemble model that combines three distinct river layers producing a final ensemble model that reflects suitable baseline environmental conditions for the species.

## Materials and methods

### Study organism and study area

Our study area (modeling region; Fig 1) comprised an approximate area of 64,750 km^2^ with 52,000 km of rivers and streams in western Oregon and was originally designated by a panel of species experts based on the current and historic distribution of foothill yellow-legged frog in Oregon (Olson and Davis 2009). The climate was cool and wet in winter with seasonal drought in the summer (17). Most of the modeling region occurred across 26 sub-basins in the Willamette Valley, Coast Range, Cascades, and Klamath Mountains (Fig 1). Our study area was primarily mountainous with upland conifer and mixed conifer-hardwood forests except for the Willamette Valley and portions of the Klamath Mountains, which contained valleys that have been greatly altered since European settlement (~1850; (12)). The Willamette Valley was characterized by agricultural development, greater disturbances to rivers, and the presence of invasive or alien species (18).

**Fig 1.**
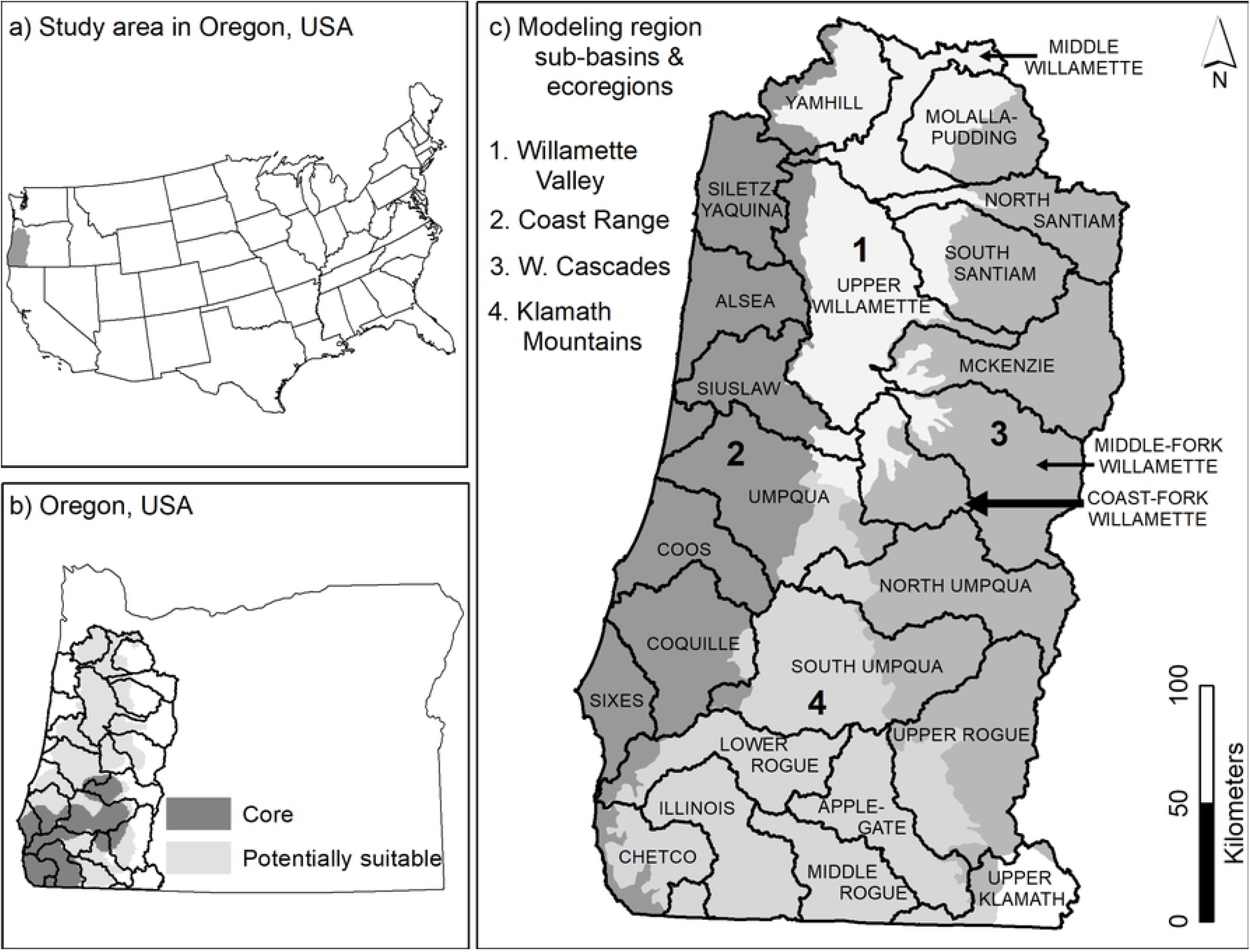
Distribution of core and potential suitable areas for foothill yellow-legged frogs in Oregon. Study area that includes distribution of core and potential areas (based on suitable baseline environmental conditions and historical species occurrence records) for yellow-legged frogs in western Oregon (a, b) spanned 26 sub-basins (c;(4)).

Foothill yellow-legged frogs primarily depend on a narrow range of in-river conditions to support all life-stages. Rivers where they occur are typically described as containing rocky, unconsolidated substrates with broad and slow-moving water, and unshaded allowing ample sunlight (2). Unconsolidated rocky substrates in slow-moving water provide stable locations for egg-laying, and seasonally warming water temperatures with ample sunlight support formation of algae, the food source of tadpoles (7,10). Breeding is seldom observed in well-shaded sites (19), and water temperature and canopy cover were the single best predictors of occurrence of tadpoles and adults in a river system in northern California (20).

### Species location data

We collated 618 locations from surveys conducted 1990 to 2019, including data used in previous modeling efforts (*n* = 237; Olson and Davis 2009); we summarize these additional data in the results. Location data were collated from a variety of surveys, including natural resource manager database (U.S. Forest Service), the Northwest Forest Plan Aquatic and Riparian

Effectiveness Monitoring Plan, and unpublished reports (7,21). All locations were within or along rivers or streams and the primary method for locating frogs were in-river surveys that located adults or egg-masses during the breeding season. As such, we assumed locations represented a portion of a locally breeding population of foothill yellow-legged frog.

To conform to the resolution of the environmental predictor layers used in each model and to reduce the likelihood that representation of environmental conditions were correlated with intensity of local sampling, location data were thinned (reduced in density) to match the resolution of our environmental layers. For example, the NorWest river layers were accurate to 1-km and we therefore limited our analysis to locations that were a minimum of 1-km from any other location. The benefit of thinning location data in this way is that using presence-only data, we cannot distinguish between survey effort and abundance of frogs such that environmental conditions in some locales could be over-represented due to higher survey effort that do not necessarily represent ‘better’ conditions for the species.

### River environmental data used in models

In-river conditions are likely to contribute most to suitable conditions for foothill yellow-legged frog, with climatic factors and upland forest cover also potentially contributing to suitable conditions (Table 1; (20)). We used three sources of data hypothesized to contribute to suitable conditions for foothill yellow-legged frog: 1) in-river variables, including stream order and flow volume, 2) climate, including precipitation and ambient temperature, and 3) composition of upland vegetation, e.g. forest cover. Foothill yellow-legged frogs are well-established as a river-obligate species, not found in perennial waterways, as such we only considered rivers as potentially providing suitable conditions that support all life stages (2).

**Table 1.** Description and predicted correlation of variables used when modeling foothill yellow-legged frog relative suitable conditions.

| Variable description | Predicted correlation | Brief description of variable prediction |
| --- | --- | --- |
| Annual radiation | + | In-season warming water supports breeding temperatures |
| Hardwood cover | + | Limited shading of rivers |
| Conifer cover | - | Greater shading relative to hardwoods |
| Total forest cover | ± | Moderate forest cover regulates water warming |
| Elevation | - | Broader, shallower rivers support breeding temperatures |
| Stream gradient | - | Low gradient limits river speed |
| Precipitation frequency | - | Summer drought provides warming profile for breeding |
| Annual precipitation | ± | Moderate precipitation to sustain rivers but limit snowpack |
| Stream order | + | Broader, shallower rivers support breeding temperatures |
| Summer max temp | + | Higher temps maintain in-season warming |
| Summer min temp | - | Higher temps maintain in-season warming |
| Summer radiation | + | In-season warming waters supports breeding temperatures |
| Winter min temp | - | Mild winters may facilitate non-breeding movements |
| Annual stream flow (CMS) | + | Larger river systems support broad channels |
| Base-flow index | - | Groundwater keeps rivers colder limiting warming water |
| Mean stream temp | + | Higher August mean temperature supports tadpole development |
| Variation stream temp | - | Stable water temperature supports breeding |
| Cumulative drainage | + | Broader, shallower rivers support breeding temperatures |

We considered three different river layers as the basis for our model with each river layer providing some unique environmental data. The extent and location of river channels, however, differed by river layer. We attempted multiple methods to combine river layers (and thus the unique information contained by each river layer) *a priori,* including buffering each river layer such that they overlapped sufficiently to provide a unified layer but could not find a method to retain resolution and accuracy of each layer. We, therefore, produced a different model for each river layer using the data available for each, but with consistent forest cover data (22), and reconciled the models into an ensemble model by combining model output from each model (explained at end of Modeling steps).

Our first model was built upon Olson and Davis (4) by using similar variables with some modification of scale (via focal radii centered on each pixel) or data source. We refer to this model as the Conservation Assessment model and it was a combination of National Hydrography Dataset (NHD) and local knowledge of river systems.

Our second model used NorWest which included modeled in-river temperature calibrated from river temperature measurements across the western United States (23). In addition to in-river temperature, NorWest estimated base-flow index, a measure of the contribution of ground-water; rivers with a greater contribution are hypothesized to be colder (24). Data for the NorWest model were available 1993 – 2015 at a resolution of 1-km river segments. We used average August river temperature and variation (1 standard error) across those years as variables in our NorWest model.

Our third model used NetMap data, which uses a variety of geo-physical models to map river characteristics. This model provided a variety of flow (cubic meters per second), stream order (highest contributing variable from previous modeling), and physical characteristics of rivers (depth, width). Stream order was highly correlated with modeled width and depth of river channel (S1 Fig). The river network produced in this layer was spatially extensive such that after determining no frog locations were within small or potentially ephemeral NetMap 1^st^ order streams and ~1% of locations were in 2^nd^ order streams and rivers, we eliminated 1^st^ order rivers from the network.

Riparian vegetation variables were produced from Gradient Nearest Neighbor forest structure and species composition imputation that used regional forest inventory plots and Landsat imagery (https://lemma.forestry.oregonstate.edu/) (25). We used 2005 as our reference year (the year from which we pulled Gradient Nearest Neighbor data) and used 30-m resolution for both NorWest and NetMap modeling.

### Modeling steps

We used five stages in modeling: 1) we examined the highest-performing scale for upland variables, 2) we eliminated highly correlated variables, 3) we applied a variable reduction procedure to produce a final model with the fewest number of variables that had similar performance metrics to a full model, 4) model calibration to prevent model overfitting and achieve the best test statistics, and 5) we produced a final ensemble model by combining all three models (Conservation Assessment, NorWest, and NetMap).

First, because biological phenomenon can be scale-dependent, we examined multiple focal radii, centered on individual pixels, for forest cover variables (30-m, 60-m, 120-m, and 240-m), retaining the radius that provided the greatest difference between species presence location data and random available locations (within our river network) that also limited variance with preference for the smallest radius.

Second, we eliminated variables that were moderately to highly correlated (pairwise Pearson correlation coefficient (r) > 0.7; e.g., S1 Fig), retaining the variable that performed best by interpreting separation between randomly generated locations (*n* = 2,000) and presence locations using density plots (S2 Fig). Although multicollinearity may be less of a problem with machine learning models compared to statistical models (26), interpretation of results between species presence and environmental variables can be problematic if variables are highly correlated (27,28). Finally, after eliminating collinear variables, we fit models using program Maxent. Maxent uses machine learning to differentiate the range of conditions where species presence location data occur from a ‘background’ sample representing available conditions for a species within the modeling region (see S3 Appendix for details on our settings). We assumed that these background locations appropriately represented conditions present in our modeling region, and therefore, interpreted logistic model output from MaxEnt as relative suitable conditions (28,29). Maxent can fit multiple variable response functions (in our case: linear, product, quadratic) that represent the complex niche space that species occupy (26).

Third, we used a variable reduction procedure that by fitting models with the fewest variables, fits the best model from the variables available and simplifies replication of our results in future modeling efforts (30). Our modeling process was to start with a full model that contained all variables (after eliminating highly correlated variables) and then to iteratively remove the variable that contributed least to model fit at each iteration whereby without the variable, training gain would remain high relative to removing any alternate variable (31). We compared model performance at each step using training gain (S4 Fig) and selected the model that had statistically similar performance metrics relative to models in previous steps that contained a greater number of variables.

Fourth, we calibrated each model to avoid model overfitting to the training data (a 75% random bootstrapped selection of frog locations) and to produce the best test statistics – Spearman rank (Rs) and area under the receiver operator curve (AUC) – based on a held-out 25% random bootstrapped selection of frog locations. To achieve this we used alternative regularization multiplier (RM) settings (0.5 – 3.0, at 0.5 intervals) and compared model training gain to test gain for signs of over fitting (training gain > testing gain) and model predictive performance using the test AUC and the continuous Boyce index (CBI) proposed by (32), which is based on the Spearman rank correlation of the model’s predicted versus expected (P/E) ratio curve We classified continuous model outputs using the distribution (95% confidence intervals [*CI*]) of the P/E ratio curve to reclassify it into four relative suitability classes. Unsuitable was considered to be P/E < 1 (upper 95% *CI* of P/E < 1), marginal as 95% *CI* overlapping P/E = 1; for suitable and highly suitable, we evenly divided all P/E > 1 values and categorized the lower 50% as suitable and the upper 50% as highly suitable.

Finally, we created an ensemble model that used output from all three models produced. Each model was categorized using the P/E curves with values of 0 (unsuitable, marginal), 1 (suitable), and 2 (highly suitable). To reconcile spatial variation in river layers, we buffered suitable classes for each river layer by a 150 m distance and assigned the maximum value within that distance. We then summed values of the three models at this resolution then resampled to 100-m pixels using a majority filter. Final output of the ensemble model ranged 0 to 6 with 6 indicating highest confidence of suitable habitat in that all models indicated highly suitable at that location.

### Human-caused disturbances to rivers

The effects of water impoundments, splash dams, and agriculture may not be entirely localized, e.g. splash dams had extensive downstream impacts and agricultural chemicals can drift and runoff into rivers (1,33). Splash dams retained and periodically releasing huge volumes of water down river and these ‘sustained pulses’ caused massive scouring of river substrates as well as heavily altering flow regimes, including increasing both the frequency and magnitude of flooding. Agricultural development can alter in-river conditions (morphology, flow) and potentially expose frogs to a variety of agricultural chemicals potentially broadcast by wind across broad distances (5,6). For these reasons and others, we averaged extent of each human-caused alteration within a 5-km radius moving window (approximately the size of a sub-watershed in our modeling region). We then created an index by re-scaling averaged extent 0 to 100.

Finally, to assess whether human-caused disturbances to rivers could have contributed to range contractions in the 20^th^ century, we compared conditions where foothill yellow-legged frogs were present or absent during contemporary surveys of locations where they were historically found (museum specimens; (7)). If conditions differed between locations where they were relocated (present) or not relocated (absent), we considered this circumstantial evidence that alterations to rivers may have contributed to extirpations. Although not definitive, we assumed this could warrant further investigation.

We summarize results using sub-basins (hydrologic unit code 8) which are, on average, 2614.3 ± 1068.5 km^2^ (mean ± 1 standard deviation; *n* = 28) and sub-watershed (hydrologic unit code 12; 77.9 ± 24.3 km^2^; *n* = 870) to summarize models. We summarized the average or 1 (highly suitable, suitable) and 0 (unsuitable) suitability values within sub-watersheds. We visually display potential threats and contemporary frog location data to illustrate the potential for threats to constrain current distribution relative to historic.

## Results

Location data included an additional 190 locations collected since 2006 (temporal extent of the Olson and Davis (2009) modeling effort), 74 of which were >1 km from a previous location and >1 km from the nearest adjacent location found after 2006. Foothill yellow-legged frog locations have been found in an additional 25 (of 118 total) sub-watersheds representing 27% of sub-watersheds with known locations. The only sub-basin that did not have a previous detection was the Umpqua, at the northernmost edge of the core of the species’ distribution in Oregon. We used the original 237 locations from Olson and Davis (2009) for the Conservation Assessment model, 290 locations for the NorWest model (1-km resolution), and 379 locations for the NetMap model (420-m resolution).

For upland forest cover, conifer cover at 60-m and hardwood cover at 240-m had the highest differentiation when comparing frog locations to random locations. We removed several variables from consideration prior to building models due to high collinearity. For NetMap, we removed river depth and width as both were correlated with stream order, and elevation in favor of winter minimum temperature (S1 Fig). Six variables used in the original Olson and Davis (4) model were highly correlated such that the updated Conservation Assessment model contained seven variables (Table 2).

**Table 2.**
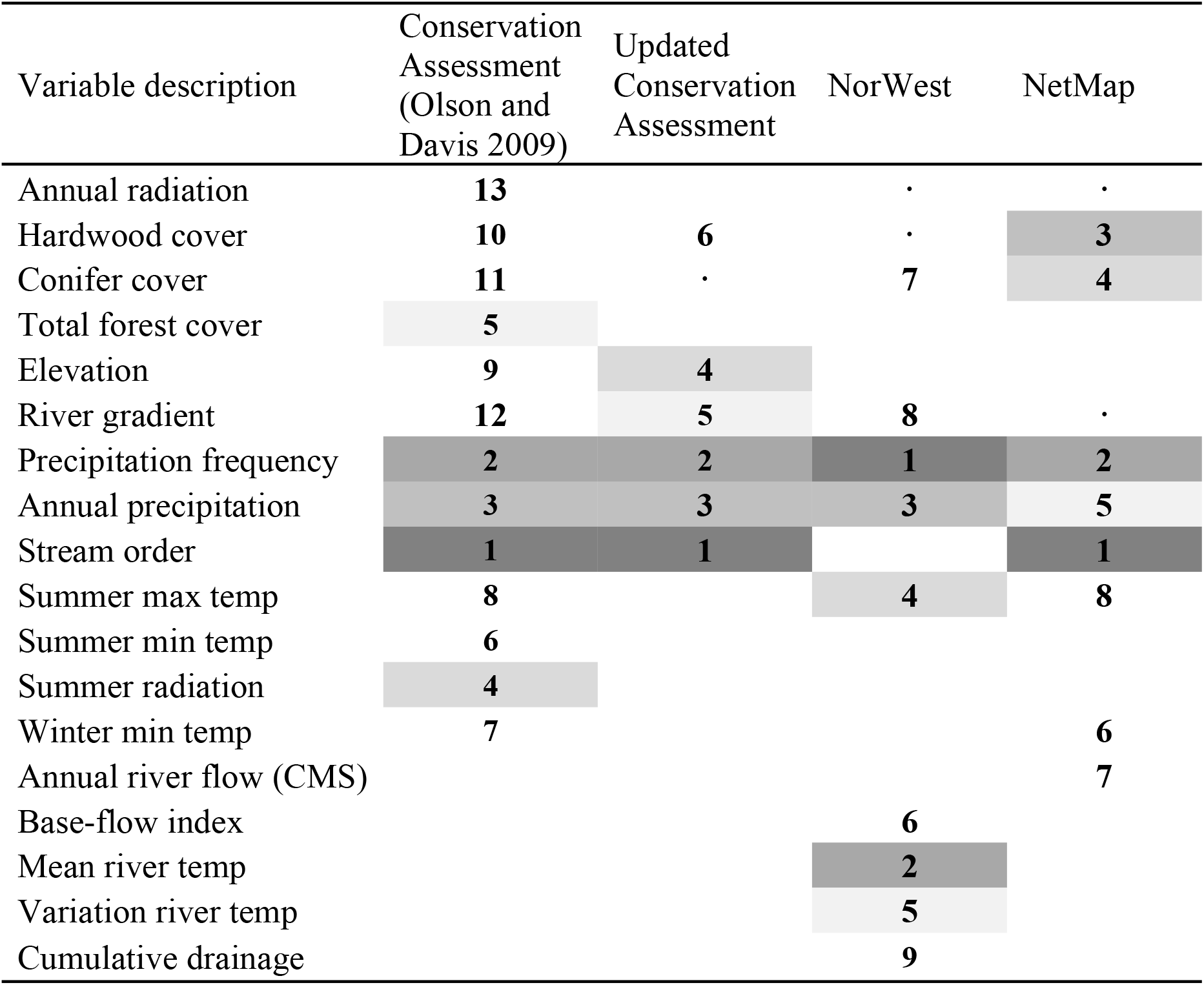
Variables included in each model and the order in which each variable was ranked. Higher ranked variables were removed later, such that 1 is the highest rank (last variable in models).

| Variable description | Conservation Assessment (Olson and Davis 2009) | Updated Conservation Assessment | NorWest | NetMap |
| --- | --- | --- | --- | --- |
| Annual radiation | 13 |  | . | . |
| Hardwood cover | 10 | 6 | . | 3 |
| Conifer cover | 11 | . | 7 | 4 |
| Total forest cover | 5 |  |  |  |
| Elevation | 9 | 4 |  |  |
| River gradient | 12 | 5 | 8 | . |
| Precipitation frequency | 2 | 2 | 1 | 2 |
| Annual precipitation | 3 | 3 | 3 | 5 |
| Stream order | 1 | 1 |  | 1 |
| Summer max temp | 8 |  | 4 | 8 |
| Summer min temp | 6 |  |  |  |
| Summer radiation | 4 |  |  |  |
| Winter min temp | 7 |  |  | 6 |
| Annual river flow (CMS) |  |  |  | 7 |
| Base-flow index |  |  | 6 |  |
| Mean river temp |  |  | 2 |  |
| Variation river temp |  |  | 5 |  |
| Cumulative drainage |  |  | 9 |  |

Model performance and final variables within each model differed with some overlap of final model variables (Table 2). The updated Conservation Assessment model had a CBI of 0.97 ± 0.02, AUC of 0.97 ± 0.01, and was modeled at regularization multiplier of 1.0; predicted suitable threshold began at 0.18. Our NorWest model at a regularization multiplier of 2.0, a CBI of 0.95 ± 0.02 and test AUC was 0.90 ± 0.01; predicted suitable began at 0.34 (Fig 2). The final NorWest model contained nine variables with precipitation frequency, mean river temperature, and annual precipitation ranked highest (Table 2). Our final NetMap model contained 8 variables, was calibrated at a regularization multiplier of 2.0, CBI of 0.97 ± 0.02, and test AUC was 0.94 ± 0.01; predicted suitable began at a value of 0.19. Stream order and precipitation frequency were the two highest ranked model variables followed by hardwood cover. Overall, stream order (when present) and precipitation frequency (frequency and annual total) ranked highly in models and final models contained at least one forest cover and ambient temperature variable (Table 2, S5 Table).

**Fig 2.**
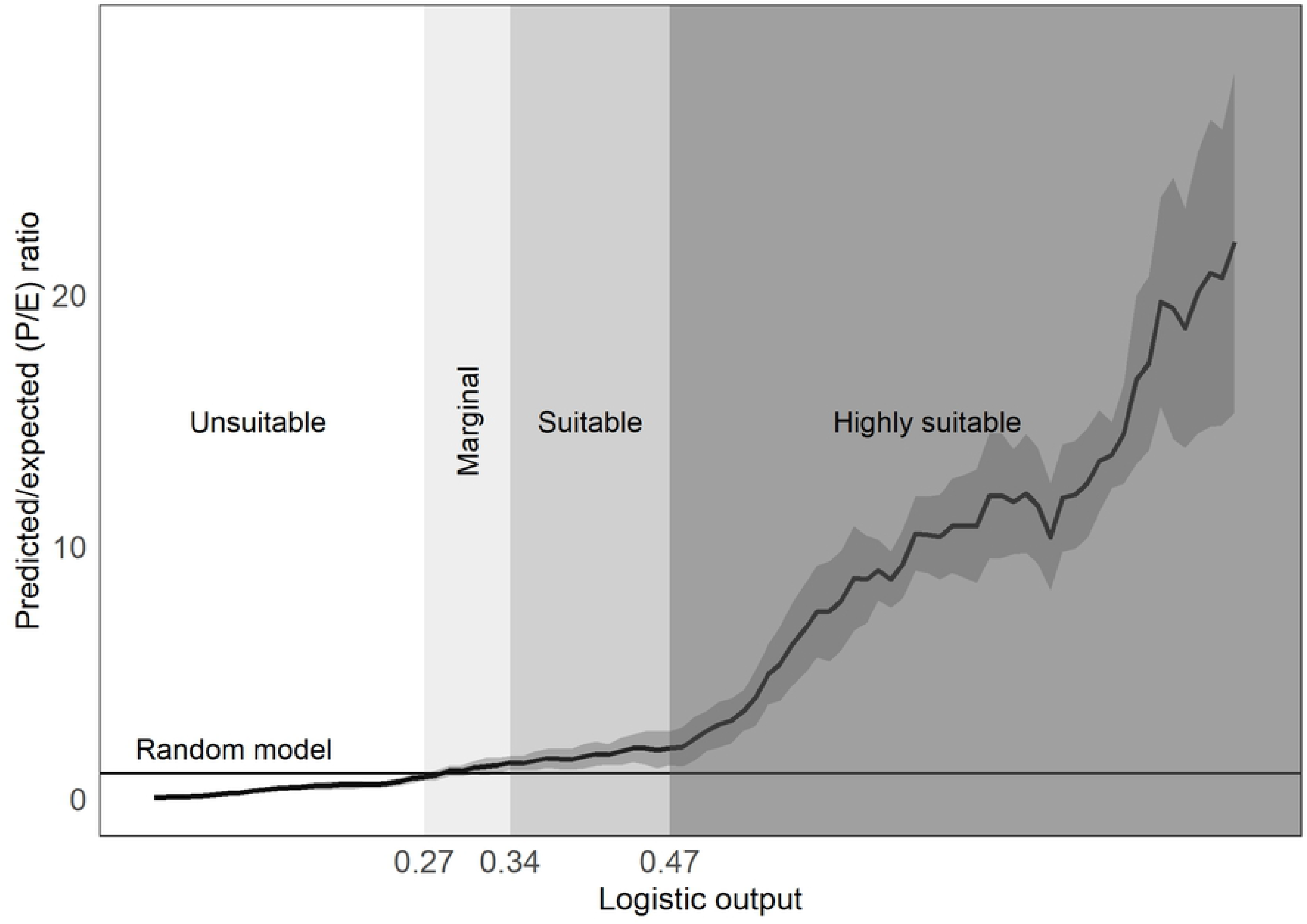
Predicted to expected ratio curve for the NorWest model. We identified breaks in relative suitability of rivers for foothill yellow-legged frogs using the 95% confidence interval, whereby at P/E < 1 (unsuitable), overlapping 1 (95% CI, marginal), and lower 50% of area at P/E > 1 (suitable), and upper 50% of area at P/E > 1 (highly suitable).

Most variables had similar response curves across models, and were broadly consistent with predicted correlation direction with some exceptions (Table 1, Fig 3, S2 Fig). Precipitation frequency (all models) and stream order (NetMap, Conservation Assessment models) had positive quadratic responses, and were inconsistent with our predictions in that we predicted a simple positive correlations for both. Mean river temperature in the NorWest model was positively correlated and annual precipitation (all models), consistent with our prediction, were positively correlated.

**Fig 3.**
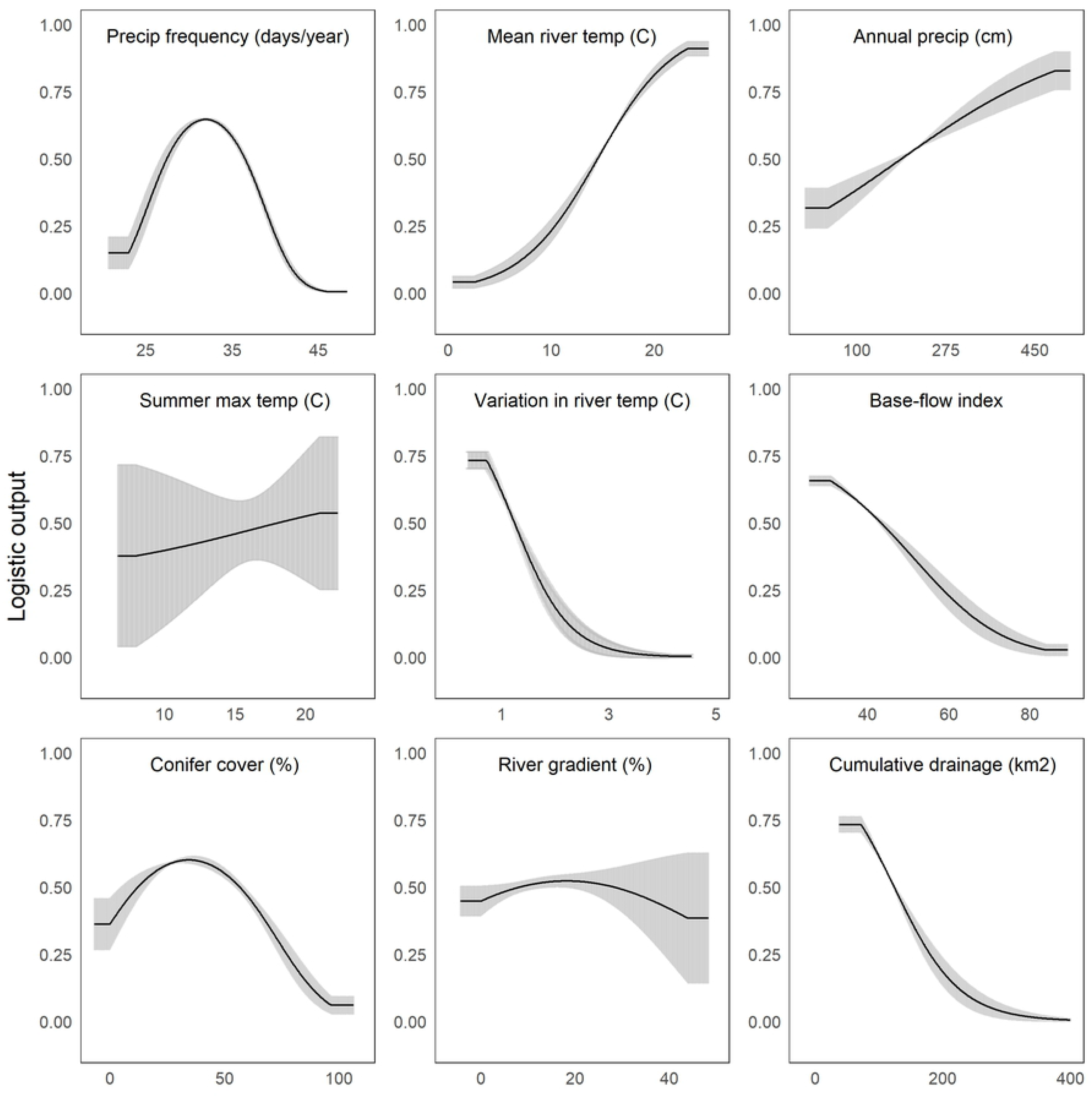
Logistic response curves for NorWest model. Response curves of variables showing the mean and variation from 10 bootstrapped replicates of the NorWest model for foothill yellow-legged frogs in Oregon. Variables are ordered by removal rank from our variable reduction procedure, with highest ranked (removed latest in procedure) at top right.

Overlap of all models appeared to be highest within the South Umpqua sub-basin. Most high-ranking sub-basins were within the Klamath Mountains and Coast Range (Figs 1, 4, 5).

**Fig 4.**
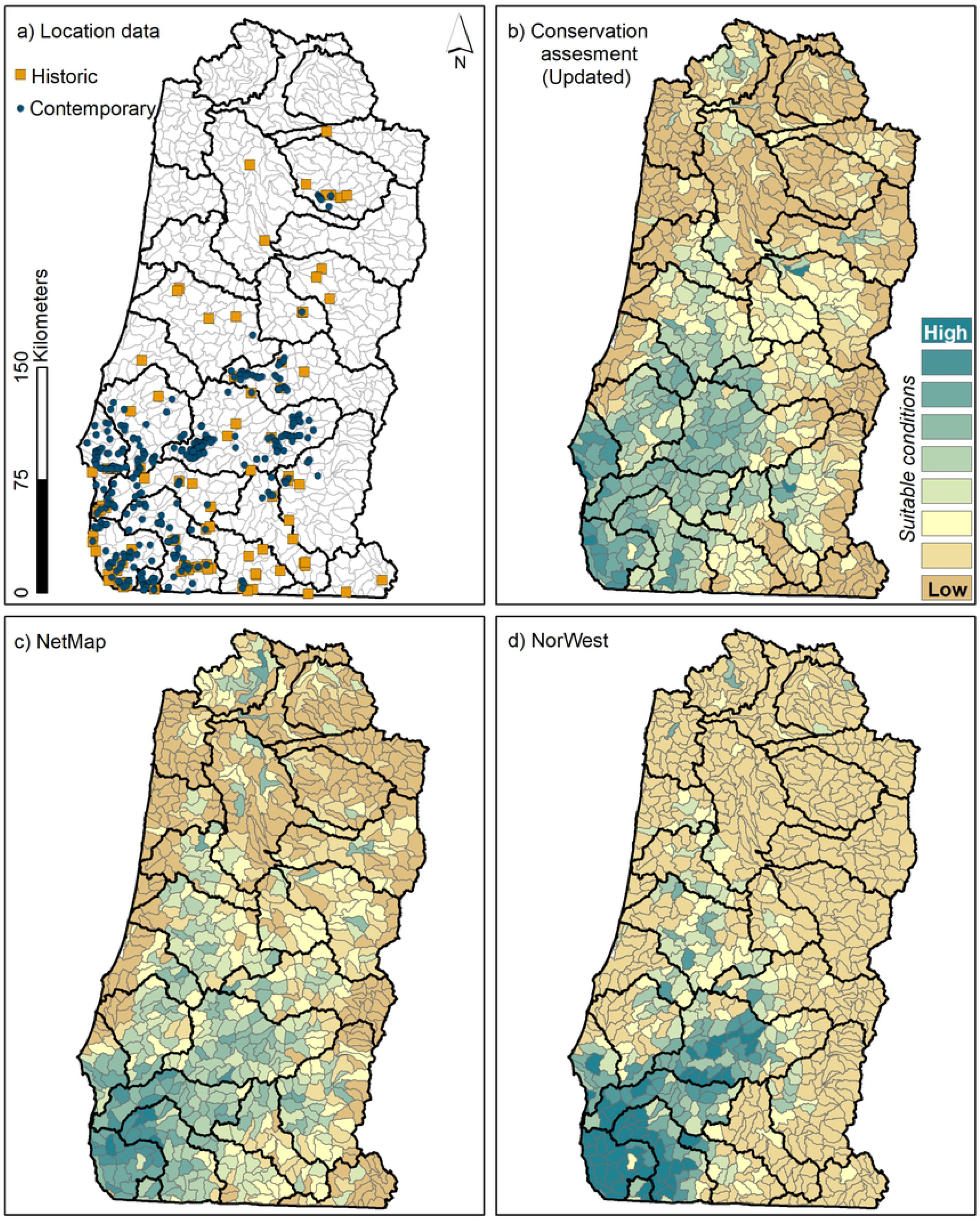
Distribution of foothill yellow-legged frog suitability using three models. Suitability is summarized by sub-watershed (hydrological unit code 12; shown in gray outline) with larger sub-basins (hydrological unit code 8) shown. Suitable conditions were summarized as the proportion of suitable or highly suitable river pixels in each sub-watershed and are represented as an indexed ranking of predicted habitat with high rankings depicted in dark blue and low in brown/tan. Panel (a) is historic and contemporary location data. Panels b – d were models produced using program Maxent, contemporary location data, and three different river layers: b) Olson and Davis 2009, c) NetMap, and d) NorWest produced from NHD-plus (USGS).

**Fig 5.**
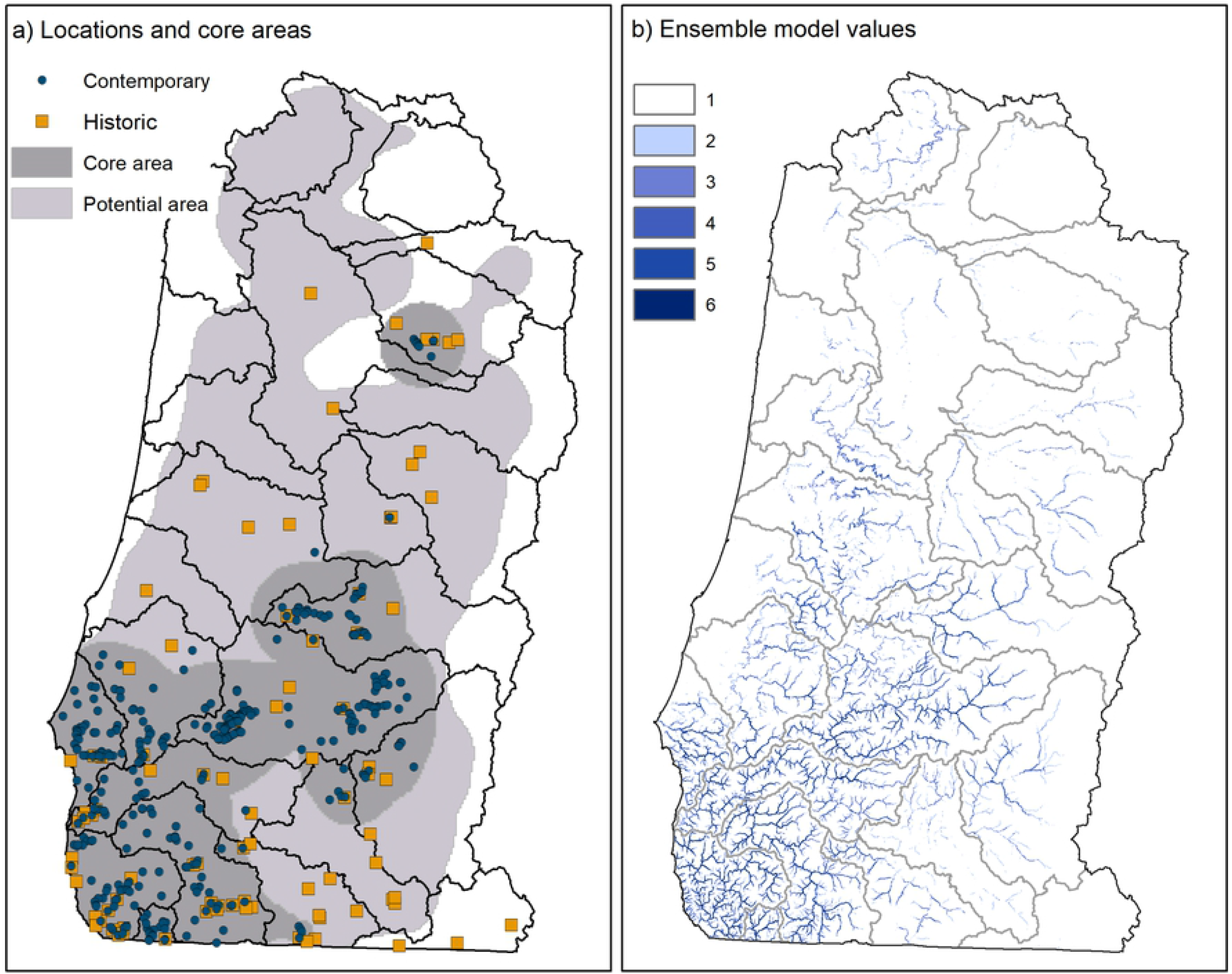
Ensemble model containing contemporary and historic locations of foothill yellow-legged frogs in western Oregon. Panel (a) shows core (dark gray) potentially suitable areas of the contemporary distribution (light gray; (4)). Panel (b) shows a multimodel ensemble whereby three values (0 = unsuitable, 1 = suitable, 2 = suitable) were summed across three models.

River disturbances were most highly concentrated in the northern and SE portion of the modeling region (Fig 6). Index values for agricultural development and splash dams were higher where frogs were absent during re-surveys of historical locations (*n* = 90; Fig 7). Splash dams primarily occurred in the forested Coast Range (Fig 6a, b) and most agriculture occurred in the Willamette Valley (Fig 1; Fig 6c).

**Fig 6.**
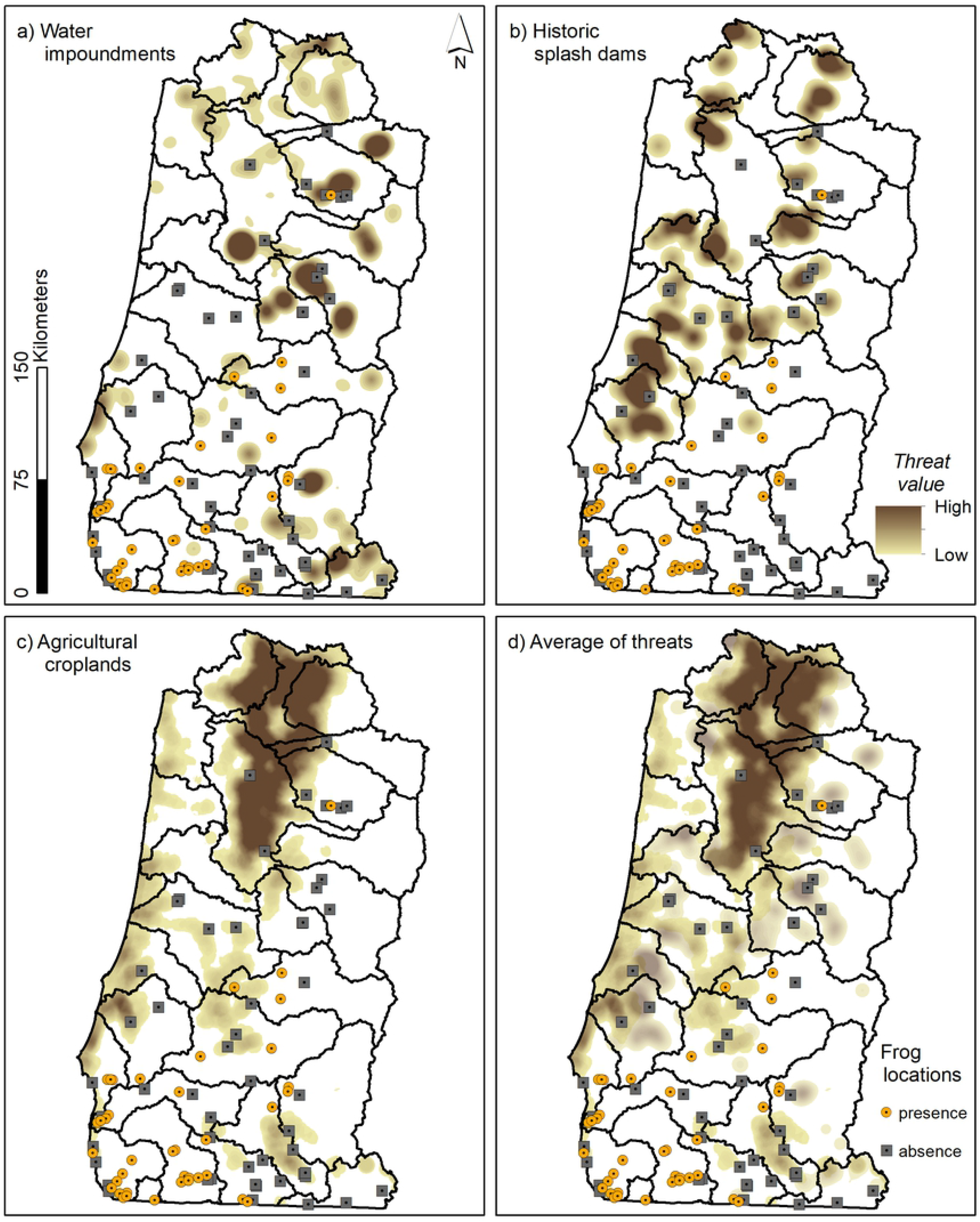
Human-caused disturbances to rivers in western Oregon. Disturbances include water impoundments ((a); e.g. large dams), splash dams on rivers used for transporting logs, agricultural croplands (c), and all threats combined with locations of foothill yellow-legged frogs overlaid.

**Fig 7.**
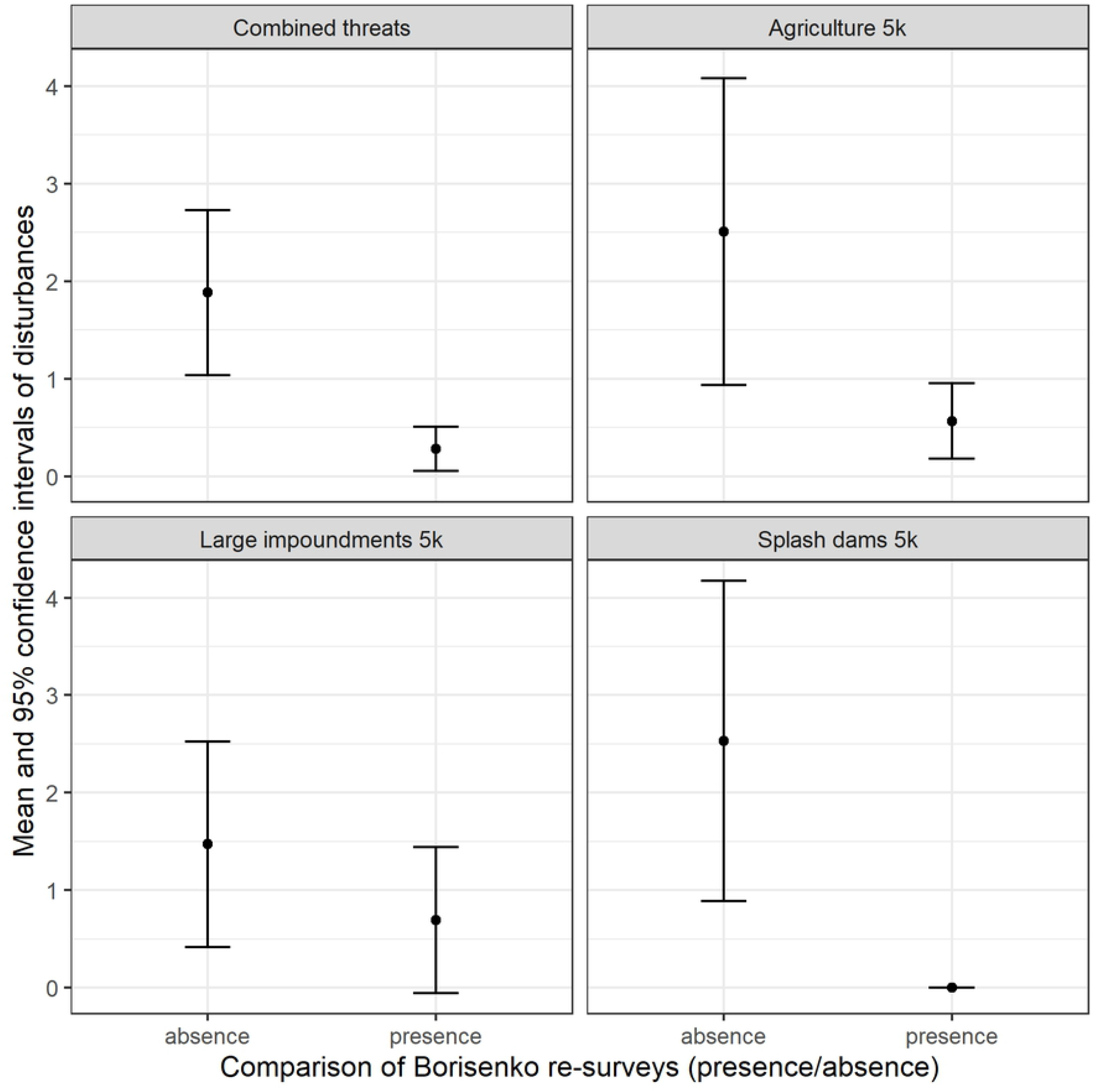
Comparison of indices of human-caused disturbances to rivers in Oregon. We used results of contemporary re-surveys of historical (museum specimens) locations whereby frogs were found (present) or not found (absence) during re-surveys. Indices were scaled 0 to 100 based on extent of human-caused alterations within a 5-km moving window which approximates the size of a sub-watershed in our modeling region. We produced indices of agricultural croplands (Agriculture 5k), large impoundments (e.g. dams; Large impoundments 5k), and timber splash-dams (Splash dams 5k).

## Discussion

Our ensemble model provided a reasoned estimation of baseline environmental conditions that explained the historic distribution of the foothill yellow legged frog in Oregon and is validated by museum records. This model and the resulting map indicated that environmentally suitable conditions were limited at the distributional edge where historical museum locations were recorded (Fig. 5a). Human-caused disturbances that are known to cause disruptions and alterations to rivers, including log-drives, splash dams, and agricultural development were most prevalent in the portions of the historic species distribution where recent surveys have failed to relocate this species. Whether alterations to rivers were of sufficient magnitude and extent to have extirpated foothill yellow-legged frogs from portions of their estimated historic distribution, including the northernmost edge, is difficult to ascertain as most disturbances occurred where frogs were thought uncommon historically (34). However, it is exactly in locations of low species abundance and distribution, that metapopulations are most at risk of extirpations attributable to localized disturbances that cause local extinctions to exceed local colonizations (35). Nonetheless, watersheds with higher levels of human-caused disruptions to the river ecosystems coincided with where foothill yellow-legged frogs are now thought to be absent (Fig 4; (4,7)).

Human-caused disturbances to rivers in Oregon can be broadly characterized as ongoing (large impoundments, agriculture) or historical (e.g. splash dams). Ongoing disturbances to rivers negatively affect foothill yellow-legged frogs, with strong evidence from water impoundments, although the species was still found in the vicinity of water impoundments during re-surveys in Oregon with little difference between index values where frogs were present or absent (Fig 7; (1,9)). Determining whether historically scoured and channelized rivers via splash dams remain unsuitable or are simply unable to be recolonized due to scarcity of local populations can inform whether efforts to restore in-river conditions are likely to succeed in re-establishing populations at the range margins.

Additional surveys for foothill yellow-legged frogs, particularly if designed to occur in watersheds with disturbed rivers adjacent to recent frog presence, could help disentangle the lasting effects of historical disturbances. As our study only identified the extent of disturbances to rivers, quantifying the magnitude of effect from disturbances done in conjunction or prior to additional surveys could provide evidence as to whether extirpations were likely to be the legacy of historical disturbances. Identifying the time periods in which legacy versus ongoing disturbances occur to natural systems can potentially inform whether populations can potentially re-establish in disturbed rivers.

Assembling multiple species distribution models into an ensemble model can provide greater confidence in understanding a species environmental relationships and inform conservation and management, but also can provide insights into uncertainty of predictions (16). By standardizing scale across models using breaks defined by the P/E curve, models used in our ensemble contributed equally (e.g. NetMap suitable > 0.19 whereas NorWest suitable > 0.34). Although we only used three breaks, more breaks could be used if a wider range of values within the ensemble model was desired, i.e. to provide a wider range of uncertainty in prediction (16). Our multimodel ensembles retained unique variables from multiple river layers, obviating the need to select a final model, and when classified using the P/E curve provided a standardized scale for comparison across models that can be used to assess uncertainty in prediction.

In-river variables contributed most to individual models and stream order provided a broad index of river conditions within our modeling region. We caution that stream order does not indicate presence of fine-scale habitat features *per se* (e.g. coarse rocky substrates) that foothill yellow-legged frogs require for breeding. Surveys of fine-scale habitat features can complement our models to target surveys within sub-watersheds that were highly ranked across models as our models likely overestimated suitable conditions and require additional ground-truthing (4,36).

Invasive species pose threats to many amphibians but interactions between foothill yellow-legged frogs and invasive species are less clear. For example, bullfrogs (*Lithobates catesbeianus*) have been posited to be a main factor leading to the declines of several ranid species in the western United States (37) but they primarily occur in lentic systems (slow-moving water) in the Willamette Valley ecoregion which may limit direct interactions with foothill yellow-legged frogs (but see (1) for a case study where indirect interactions, including bullfrogs, dams, and disease may be correlated with declines of foothill yellow-legged frogs). Although we considered invasive species distributions as a potential disturbance, invasives may simply be symptomatic of other human-caused disturbances to river hydrology and thus, already represented within those that we considered.

We anticipate the broadest changes to suitable conditions for foothill yellow-legged frogs occur outside of our model variables. Specifically, our strongest model variables were river geomorphology or network topology (e.g. stream order), and annual precipitation which are not expected to change much through time, whereas disturbances to river hydrological processes have been frequently identified to negatively affect the species, were not directly represented in our models, and could potentially fairly rapidly alter in-river conditions (9). When model variables are not the drivers of changes to contemporary species distribution, future modeling efforts may benefit from directly modeling potential disturbances, including magnitude of effect, to the species.

Disturbances can frequently limit species at their range margins but whether disturbance history in this area was simply correlated with lack of species occurrences or causative is unknown. Foothill yellow-legged frogs appear to be sensitive to a broad range of human-caused disruptions to rivers in Oregon, perhaps indicative of their narrow ecological tolerance to in-river conditions. Additional surveys and analysis, including assessing magnitude of river disturbances, could inform whether frogs can potentially re-establish where they have been extirpated and that have experienced high levels of human-caused disturbances and alterations.

## Acknowledgments

We would like to thank Kelli van Norman, Jeffrey Dillon, and Rob Huff for their insightful comments on drafts of the manuscript. We would also like to thank xx reviewers for contributions to the manuscript. We received funding and support from the Pacific Northwest Research station and the Interagency Special Status / Sensitive Species Program of the USDI Bureau of Land management, USDA Forest Service region 6.

